# A deep learning approach reveals unexplored landscape of viral expression in cancer

**DOI:** 10.1101/2022.06.26.497658

**Authors:** Abdurrahman Elbasir, Ying Ye, Daniel E. Schäffer, Xue Hao, Jayamanna Wickramasinghe, Konstantinos Tsingas, Paul M. Lieberman, Qi Long, Quaid Morris, Rugang Zhang, Alejandro A. Schäffer, Noam Auslander

## Abstract

About 15% of human cancer cases are attributed to viral infections. To date, virus expression in tumor tissues has been mostly studied by aligning tumor RNA sequencing reads to databases of known viruses. To allow identification of divergent viruses and rapid characterization of the tumor virome, we develop viRNAtrap, an alignment-free pipeline to identify viral reads and assemble viral contigs. We utilize viRNAtrap, which is based on a deep learning model trained to discriminate viral RNAseq reads, to explore viral expression in cancers and apply it to 14 cancer types from The Cancer Genome Atlas (TCGA). Using viRNAtrap, we uncover expression of unexpected and divergent viruses that have not previously been implicated in cancer and disclose human endogenous viruses whose expression is associated with poor overall survival. The viRNAtrap pipeline provides a way forward to study viral infections associated with different clinical conditions.

## Introduction

Viral infections have a causal role in approximately 15% of all cancer cases worldwide^1^. Viruses linked to cancer are generally divided into direct carcinogens, which drive an oncogenic transformation through viral oncogene expression, and indirect carcinogens, which may lead to cancer through mutagenesis associated with infection and inflammation. To date, seven viruses have been classified as direct carcinogenic agents in humans^2^. Among these, the high-risk subtypes of human papillomavirus (HPV) are the causative agent of approximately 5% of human cancers. Chronic hepatitis B virus (HBV) or hepatitis C virus (HCV) infections are associated with most hepatocellular carcinoma cases. More recently, advances in sequencing technologies have contributed to better appreciation of the high burden of viral infections in cancer, exemplified by the Kaposi’s sarcoma herpesvirus and the Merkel cell polyomavirus, which were discovered based on nucleic acid subtraction to cause Kaposi’s sarcoma and Merkel cell carcinoma, respectively^2^. The discovery of oncogenic viruses, starting with the Rous sarcoma virus^3^, has been critical for understanding mechanisms driving cancer evolution and for improving cancer prevention and intervention strategies. However, the burden of viral infections in cancer is thought to remain underappreciated by much of the cancer research community^4^.

Since the advent of next-generation sequencing, new viral strains are typically identified from large-scale DNA or RNA sequencing data based on sequence similarity to known viruses. The Cancer Genome Atlas (TCGA) has become a principal resource for identification of viral sequences in cancer tissues. Several studies screened TCGA DNA sequencing data to characterize known viruses in cancers^5^, and analyze host integration sites for viruses such as HBV that integrate into the human genome^6^. Other studies used RNA sequencing to screen for known viruses in the human transcriptome^7,8,9,10^, and to discover novel viral isolates^10^. Most recently, a few studies combined DNA and RNA sequencing to quantify presence of known cancer-associated viruses in human cancers^11,12^. However, the set of sequenced viral clades and the set of viral clades known to infect humans are both incomplete. Viruses and cancers have rapidly evolving genomes, and a new cancer-associated virus may have little sequence similarity to known viruses isolated outside of the tumor micro-environment. This issue is exacerbated when analyzing short reads, which are typical to RNA sequencing technologies. Therefore, discovery of new and divergent cancer viruses remains highly challenging with existing strategies^13^. For detection of bacterial viruses from metagenomic DNA sequencing, several machine and deep learning techniques have been recently developed. These methods overcome some of the limitations associated with homology-based approaches and rapidly identify viral reads including novel and divergent viruses^14,15,16,17,18^. More recently, methods have been developed to identify viruses that have potential to cause humans infections^19,20^. These recently developed methods suggest that deep learning methods to detect viral reads from RNA sequencing have the potential to uncover novel and divergent viruses in human tissues.

Here, we develop a framework, named viRNAtrap, that employs a deep learning model to accurately distinguish viral reads from RNA sequencing, and utilizes the model scores to assemble viral contigs. We apply viRNAtrap to 14 cancer types from TCGA (selected based on potential viral relevance to oncogenesis), to perform an exploratory data analysis and characterize the landscape of viral infections in the human cancer transcriptome. We demonstrate the ability of viRNAtrap to identify different types of viruses that are expressed in tumors by constructing three viral databases and comparing viRNAtrap findings to sequences in those databases. We first evaluate known cancer-associated viruses that are expressed in different tumor types. Then, we curate a database of potentially functional human endogenous retroviruses (HERVs) and analyze expression patterns of different HERVs across human cancers to find that HERV expression is associated with poor survival rates. Finally, we employ viRNAtrap to identify divergent viruses that are expressed in tumor tissues. Notably, we identify a *Redondoviridae* member that is expressed in head and neck carcinomas, a *Siphoviridae* member that is expressed in 10% of high grade serous ovarian cancers, and a *Betairidovirinae* member that is expressed in more than 25% of endometrial cancer samples. In summary, we present the first deep learning-based method to identify viruses from human RNA sequencing and demonstrate its ability to rapidly characterize viruses that are expressed in tumors and uncover viral instances that have not been previously found in these samples using alignment-based methods. viRNAtrap can be applied to identify new viruses that are expressed in a variety of other malignancies, introducing new avenues to study viral diseases.

## Results

### The viRNAtrap framework

To identify viruses in the human transcriptome, we first trained a neural network to distinguish viral reads based on short sequences. We collected positive (viral) and negative (human) transcripts that were segmented into 48bp fragments and divided into training and test sets (Figure 1a, Methods). We used different metrics to evaluate the ability of the model to identify viral sequences based on short segments. The model yielded test-set performance: area under the receiver operating characteristic curve (AUROC) of 0.81, area under the precision recall curve (AUPRC) of 0.82 (Figure 1b), accuracy of 0.71, recall of 0.83, precision of 0.67 and F1-score of 0.74 (Figure 1c). We compared the performance of this model to previous models trained to identify viruses, namely DeepViFi^16^, DeepVirFinder^15^, ViraMiner^21^, as well as a method called ‘off-the-shelf Seq2Seq’ compared through DeepViFi^16^, that does not use much domain-specific knowledge about viruses (Methods). Importantly, our model outperformed other methods in all measures, except for precision, for which DeepVirFinder outperformed all other methods (Figure 1b-c). However, precision is less critical for this framework because alignment steps are used to further filter out negatives. Importantly, DeepViFi^16^, DeepVirFinder^15^, and ViraMiner^21^ were previously not trained or evaluated for RNA sequencing or 48bp reads, which is likely the reason that these methods are less appropriate in that context without specific optimization (see Methods). Examining the average model performance across segments from different human viruses, we find that human singlestranded DNA viruses from taxon *Monodnaviria* were assigned with high confidence, whereas, for RNA viruses, we observed more variation in model confidence. For example, the model confidently predicted the viral origin of sequences from Ebola and influenza viruses but assigned borderline scores to sequences from several *Phenuiviridae* members such as *Dabie bandavirus* (Figure 1d, Supplementary Data 1).

**Figure 1.**
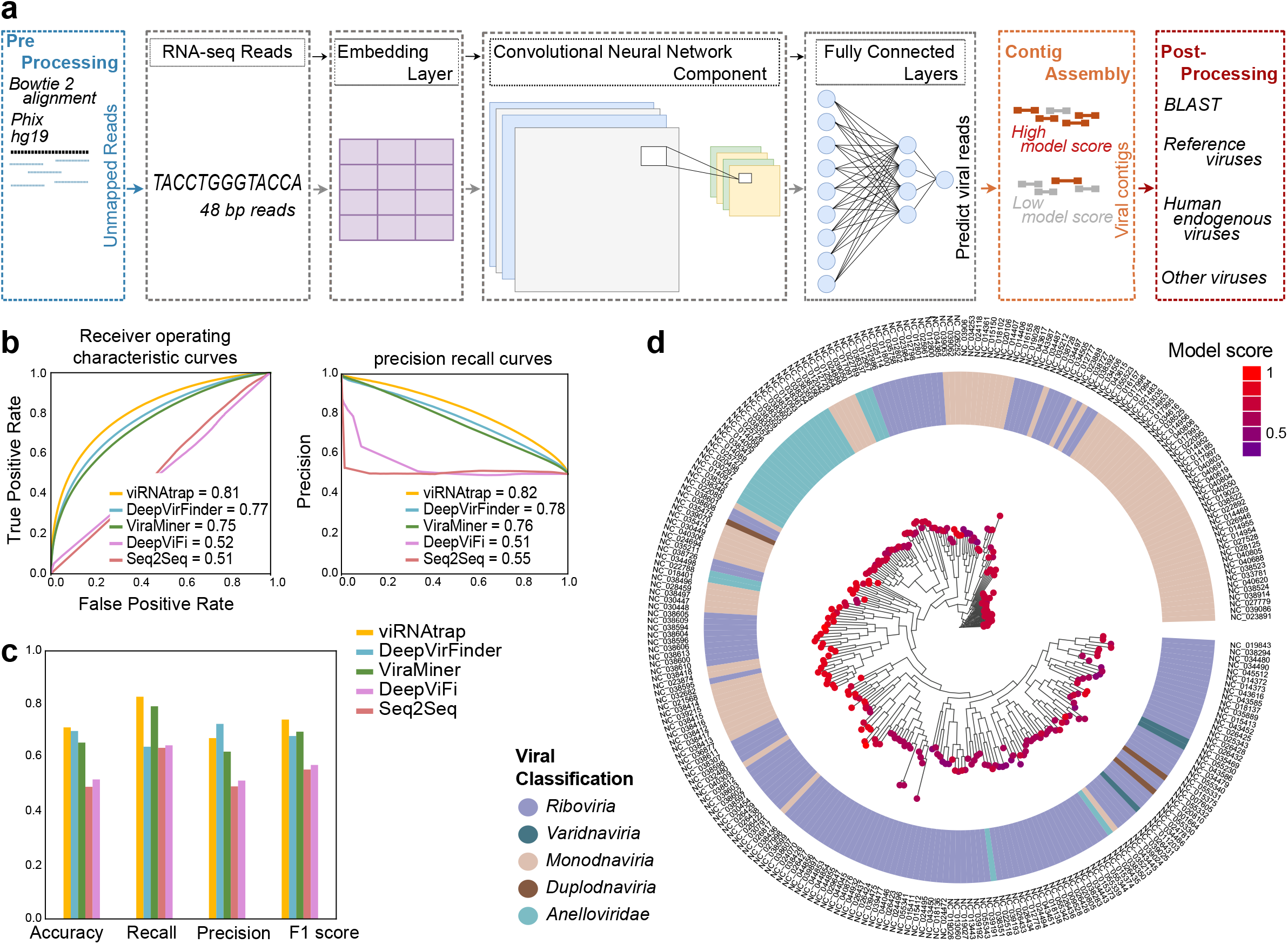
Training and evaluation of the viRNAtrap framework. **(a)** A schematic overview of the viRNAtrap framework. Unmapped reads were extracted and given as input to the neural network, to extract the viral reads and assemble viral contigs, that were compared against three viral databases using blastn. **(b)** Receiver-operating characteristic and precision-recall curves showing the model performance when viRNAtrap and models used for comparison were applied to the test set. **(c)** Bar plots showing different metrics to evaluate the model performance for the test set, for viRNAtrap and models used for comparison. **(d)** A phylogenetic tree showing the model scores for sequences from different human viruses with the respective virus classification (using average assigned score for each virus). Source data are provided as a Source Data file 1.

Based on the trained neural network, we built a computational framework (Figure 1a, Methods) to identify viral contigs from tumor RNAseq and applied the framework to 7272 samples from 14 cancer types in The Cancer Genome Atlas (TCGA)^22^, from which 6717 were tumor samples and 555 were non-cancer samples matched to a cancer sample from the same individual (Supplementary Data 2). In pre-processing, we extracted reads that were not aligned to the human genome (hg19) or to the phiX phage^23^ that was identified as a frequent contaminant.

The computational framework, named viRNAtrap, was then applied to unaligned RNA reads (to reduce the running time of viRNAtrap), to detect viral reads and assemble predicted viral contigs. Finally, in post-processing analysis, we used blastn^24^ to compare the assembled viral contigs to three curated viral databases. We identified viral contigs originating from reference viruses that are expected in cancer tissues, human endogenous viruses, and candidate novel or more divergent viruses, which are expressed in different cancer types

### Identifying reference tumor viruses

We first characterized the presence of known cancer-associated human viruses in different tumor types. High-risk human *Alphapapillomavirus* strains (HR-□HPVs) were most frequently detected; the type observed in the majority of TCGA samples is HPV16. This is expected because HR-□HPVs, such as HPV16 and HPV18, underlie approximately 5% of cancer cases worldwide^25^ while low-risk human *Alphapapillomavirus* (LR-□HPV) strains, such as HPV54 and HPV201, are mostly associated with the development of genital warts but not cancer^26^. We found at least one HR-□HPV in 288 CESC samples (286 squamous cell carcinoma samples and 2 non-cancer samples). We found 61 HNSC samples, and a total of 14 samples across other cancer types, that contain a contig from at least one HR-□HPV (Figure 2a). LR-□HPVs were identified in a small set of samples mostly from matched non-cancer tissues, including cervix and head and neck (Figure 2a, Supplementary Data 2,3).

**Figure 2.**
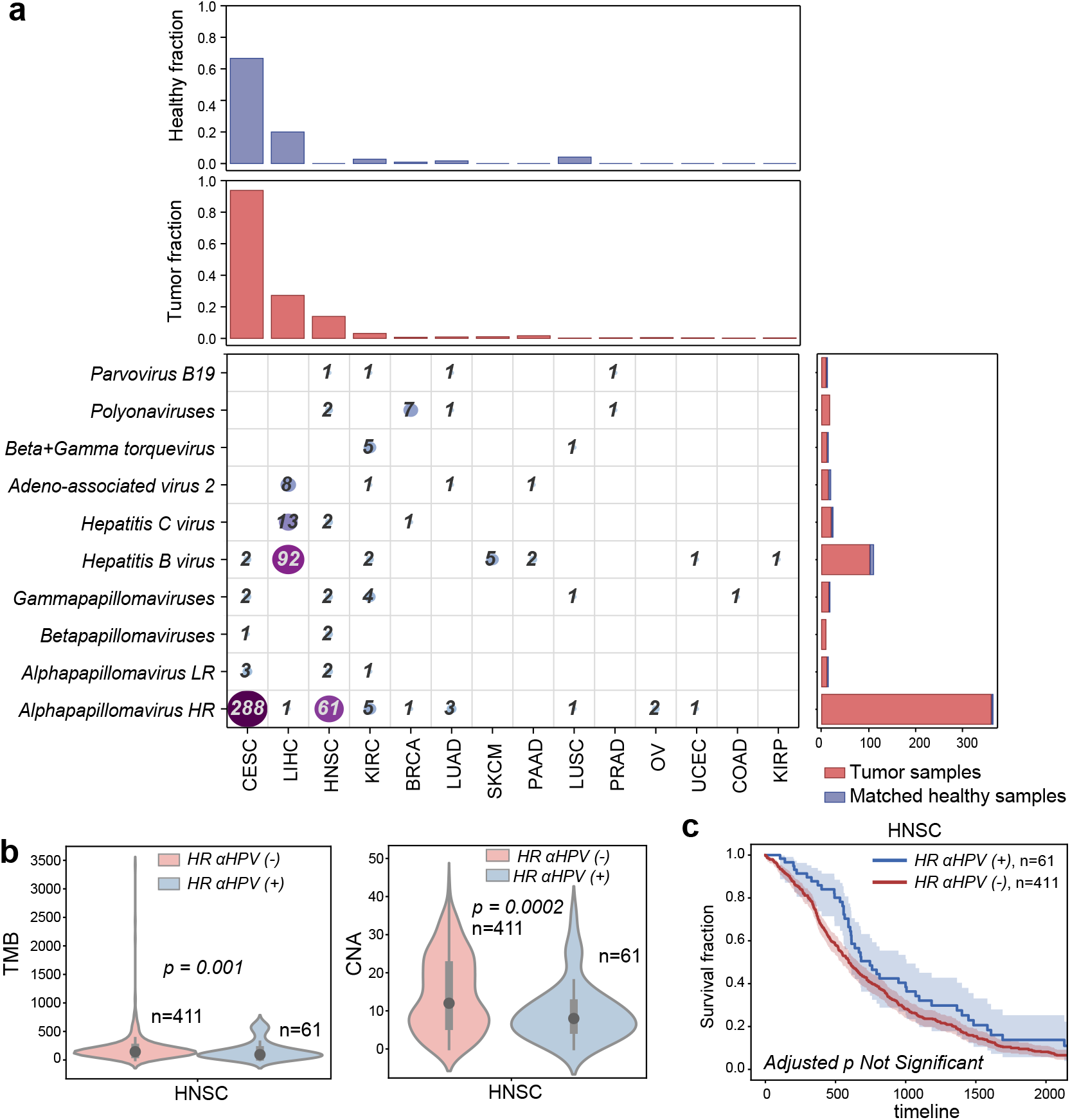
Reference human viruses expressed in different tumor types. **(a)** Heatmap showing the total number of virus-positive samples identified from RNA-sequencing in different tumor tissues. Top panels show the fraction of tumor and non-cancer samples in which viruses were identified. Right panels show the number of viruses found in tumor and non-cancer samples. **(b)** Violin plots comparing the tumor mutation burden (TMB) and the number of chromosome-level copy number alteration (CNA) between HNSC patients where expression of high-risk alpha papilloma viruses was detected vs those patients where expression of high-risk alpha papilloma viruses was not detected. Black dots represent the medians, and the boundaries of the violin plots refer to the maximum and minimum values, respectively. Two-sided Wilcoxon rank-sum p-value is reported. **(c)** Kaplan-Meier curves comparing the survival rates between HNSC patients where the expression of high-risk alpha papilloma viruses was detected (blue curve) vs those where the expression of high-risk alpha papilloma viruses was not detected (red curve). The FDR adjusted two-sided log-rank p-value is not significant (Supplementary Table 1). For Kaplan–Meier curves, shaded areas represent the confidence interval of survival. Source data are provided as a Source Data file 2.

Hepatitis B virus (HBV) is the second most frequently detected virus across TCGA samples. HBV infections and Hepatitis C virus (HCV) infections are two primary causes of liver cancer and may co-occur in a patient^11^. We found HBV expression in 85 LIHC tumor samples and 7 non-cancer samples, and HCV in 13 LIHC tumor samples. HBV was also found in a few tumor samples and matched non-cancer samples from other cancer types (Figure 2a). By comparing the samples predicted as virus-positive by vRNAtrap to the samples annotated as virus-positive in the TCGA clinical annotations, we found that the true positive rates of viRNAtrap were above 95% for HR-□HPVs (in CESC and HNSC), and for HCV and HBV in LIHC, supporting that viRNAtrap correctly identifies samples expressing known cancer viruses (Supplementary Figure 1). In addition, viRNAtrap found adeno-associated virus 2 (AAV2) in 8 LIHC samples, 6 from tumors and 2 from non-cancer samples. AAV2 is a small DNA virus that has the potential to integrate into human genes and contribute to oncogenesis, although the current evidence is insufficient for AAV2 to be included in the consensus list of oncogenic viruses^27,28^. A recent study that addressed discrepancies in AAV2 expression across TCGA samples found at least one AAV2 read in 11 LIHC samples^27^. However, in three of these samples only one AAV2 read was found, which is difficult to detect with the viRNAtrap pipeline. Notably, previous studies that systematically characterized viral presence across TCGA did not identify AAV2 in more than six LIHC samples^11,27^, demonstrating the sensitivity of viRNAtrap compared to other computational methods. We additionally detected AAV2 in one KIRC sample, one PAAD sample and one matched non-cancer sample from LUAD (Figure 2a).

We found several samples that express human polyomaviruses, especially polyomaviruses 6 and 7. Most notably, we found seven BRCA samples and two HNSC samples that express polyomaviruses. We additionally found Parvovirus B19 sequences in a few samples^29^ (three cancer and one matched non-cancer); this virus has been mostly associated with normal tissues^30^, but was also previously identified in isolated tumor cases^31,32^. We investigated possible genomic correlates of the expression of these viruses, including the tumor mutation burden (TMB, the rate of somatic mutations in a tumor, which is a biomarker and is annotated for all TCGA samples), and the chromosome-level aneuploidy (Methods). We found that HR-□HPV-positive samples have lower TMB and aneuploidy levels compared to HR-□HPV-negative samples (Figure 2b). In contrast, LIHC cancer patients positive for HBV showed significantly higher TMB compared to HBV-negative samples (Supplementary Figure 2). We additionally examined the association between the expression of known oncoviruses and overall survival. While none of the associations were significant after adjustment for multiple hypotheses (Supplementary Figure 2, Supplementary Table 1), we found a trend that HR-□HPV-positive HNSC patients have better survival compared to HR-□HPV-negative patients (by the Kaplan Meier curves Figure 2c), which is confirmatory of previous studies^33,34^. We also found a positive association between viral presence and overall survival of LIHC patients with HBV (Supplementary Figure 2, Supplementary Table 1).

### Uncovering expression patterns of HERVs in cancer tissues

To further demonstrate the utility of viRNAtrap, we analyzed the expression of HERVs across different tumor types in TCGA (HERVs were not used to train the viRNAtrap model). HERVs constitute approximately 8% of the human genome; most HERV sequences are remnants of ancestral retroviral infection that became fixed in the germline DNA^35,36^. HERV proteins are found expressed in different conditions including cancer tissues^37,38,39,40,41^. Specifically, the HERV-K family, which most recently integrated to the human genome and is one of the most abundant HERV families in the human genome (along with HERV-H), was previously reported in tumor tissues and cell lines^42,43^. Moreover, recent findings reported association between HERV expression and poor survival rates^12,36,44–46^.

To comprehensively characterize HERV members that are expressed in different tumors, we established a database of potentially functional HERVs that were extracted from the human genome (Methods). The viRNAtrap contigs were aligned against this database, to identify patterns of HERV expression in the 14 cancer types considered throughout this study.

As expected, we found that the most abundantly expressed HERV families are HERV-K and HERV-H. The fraction of samples expressing different individual HERV members was used to cluster tumor types. Interestingly, we found that squamous cell carcinomas (including cervical, lung, and head and neck) are clustered together based on the proportional distribution of expressed HERV members (Figure 3a). The HERVs that are most abundantly expressed across different cancers include some that are in proximity to cancer-associated genes or single nucleotide polymorphisms (SNPs) (Supplementary Data 3,4). Specifically, one HERV-H member (chr2:204826665-204832368) is located 365bp from the *ICOS* (Inducible T-cell costimulatory) gene, which has been associated with tumor immune responses^47,48,49,50^. In addition, one HERV9 member (chrX:150718827-150731816) is located 330bp from the *PASD1* cancer/testis antigen gene (each of these two HERVs are found in 10 TCGA samples, Supplementary Data 4,5).

**Figure 3.**
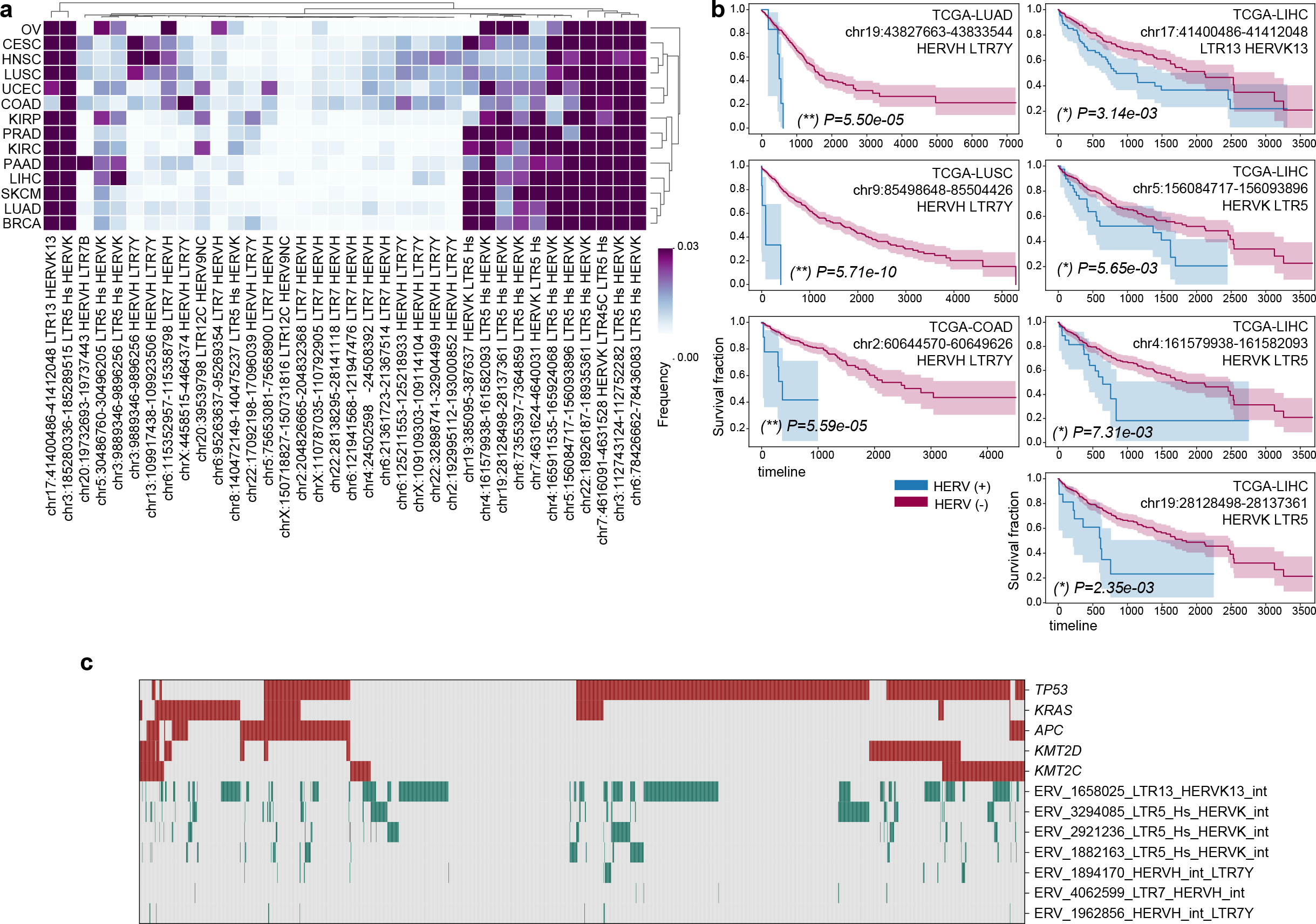
Human endogenous retroviruses (HERVs) expressed in different cancer types. **(a)** Heatmap clustogram clustering the proportion of HERVs across different tumor types. The rows are 14 TCGA tumor types. The 36 columns are the 36 distinct HERVs with the highest expression in human cancers, mapped to unique regions in the genome (Supplementary Data 5). **(b)** Selection of Kaplan-Meier curves comparing the survival rates between patients in which any HERV reads were detected (blue curves) versus those in which no HERV reads were detected (red curves). The unadjusted two-sided log rank p-values are reported. (**) global FDR q<0.05, (*) cancer-type specific FDR q<0.05. For Kaplan–Meier curves, shaded areas represent the confidence interval of survival. Additional significant associations between HERV and survival are reported in Supplementary Data 12. **(c)** Heatmap showing somatic mutations in major cancer driver genes (selected are the most frequently mutated driver genes in these samples, red) and the expression of HERVs that are significantly associated with survival in LIHC, LUAD, LUSC and COAD (green). Source data are provided as a Source Data file 3.

We investigated associations between HERV transcript presence and patients’ overall survival (Figure 3b). In agreement with previous studies^12,36,44–46^, we find that patients with HERV-K-and HERV-H-positive cancer samples have significantly lower overall survival compared to HERV-K-and HERV-H-negative patients in COAD, LUSC, LUAD and LIHC. Notably, every significant association that we identified between HERV presence and overall survival in these cancer types is negative (Figure 3b, Supplementary Table 2).

To investigate the link between HERV expression and poor survival, we compared the TMB and aneuploidy scores between patients expressing HERVs and those without HERV expression. HERVs that were associated with poor survival were not associated with TMB or aneuploidy (Supplementary Data 6). We found that HERVs associated with poor overall survival were generally more likely to be expressed in the presence of somatic mutations in frequently mutated cancer driver genes, such as *TP53, KRAS, ARID1A* and *PTEN* (using hyper-geometric enrichment, Supplementary Data 7). However, we did not find a strong association with mutations in any specific gene, and HERV expression was found even in samples with no somatic mutations in any of these genes (Figure 3c, Supplementary Data 8)

### Finding divergent viruses in human cancer

We next investigated tumor expression of divergent viruses that have rarely or never been previously reported in human cancers. We aligned the contigs produced by viRNAtrap against a database of viruses (Methods) from different hosts that were not expected to be found in tumor tissues, including human, bat, mouse, insect, plant, and bacterial viruses. (Figure 4a). We found multiple contigs of mosaic plant viruses in distinct samples from most tumor types, especially adenocarcinomas. For example, watermelon mosaic virus was found in 3 colorectal cancer samples, and Bermuda grass latent virus, which was previously reported in a COAD sample^10^, was identified in multiple samples from three cancer types (COAD, LIHC, UCEC; Figure 4a). Mosaic plant viruses have been previously detected in human faeces^51,52^, which could suggest viral entry and travel through the digestive tract. However, it is unclear how mosaic plant viruses would reach other tumor tissues, such as the liver and the endometrium, and whether these are associated with an unidentified source of laboratory contamination.

**Figure 4.**
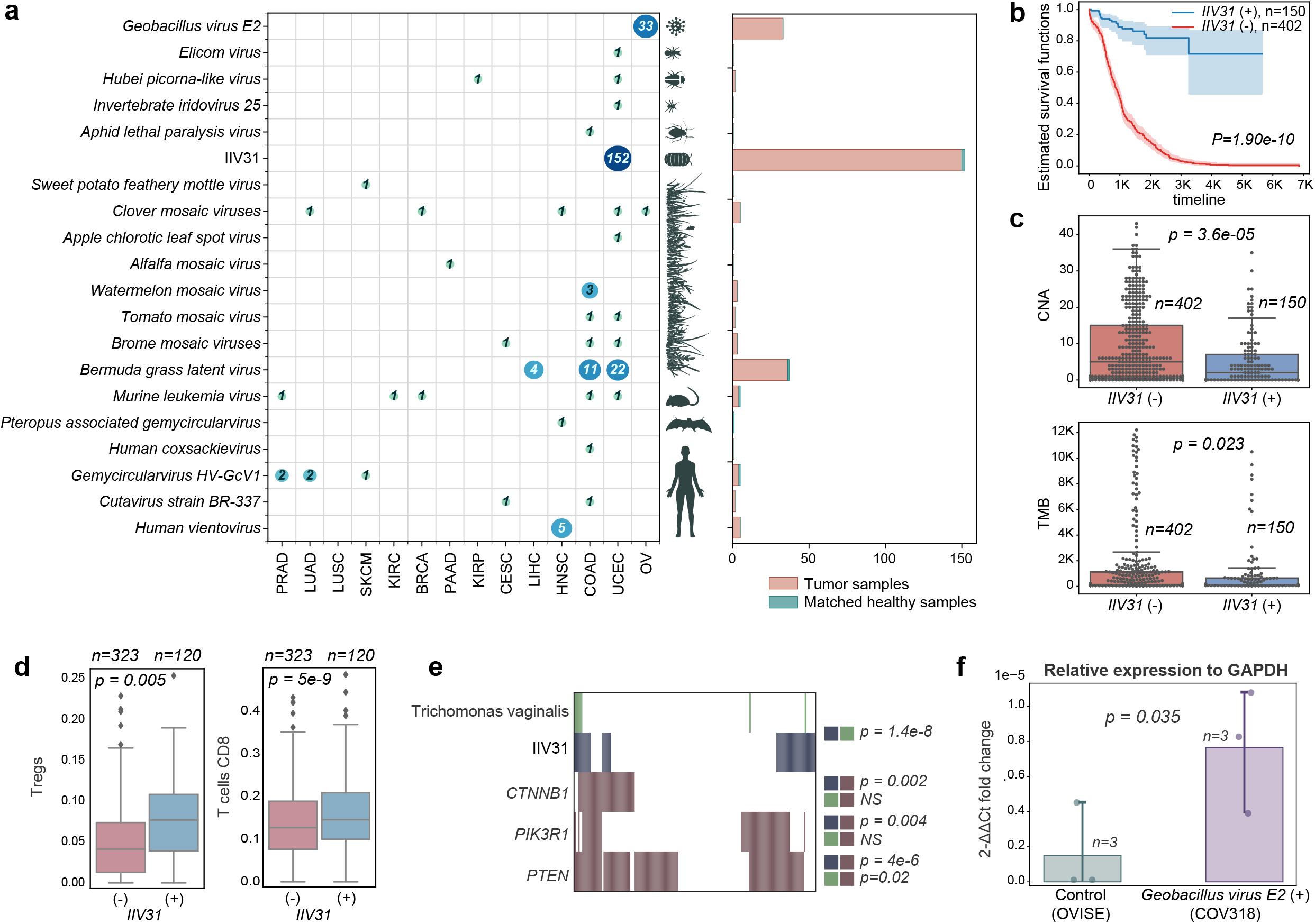
Unexpected and divergent viruses infecting different host taxa across TCGA samples. **(a)** Unexpected and divergent viruses expressed in TCGA samples. Each row in the matrix represents one virus and the entry in each column indicates the number of cancer samples of each type in which each virus was detected. The canonical hosts of each virus are depicted at the left of the matrix. At right, the aggregate number of tumor and normal samples containing reads of each virus are shown in a bar plot. **(b)** Kaplan-Meier curves comparing the survival rates between patients in which IIV31 reads were detected (blue curves) vs those where viral reads were not detected (red curves). For Kaplan–Meier curves, shaded areas represent the confidence interval of survival. The log rank p-value is reported. **(c)** Box plots comparing the chromosome-level copy number alteration (CNA, top panel) and the tumor mutation burden (TMB, bottom panel) between cancer patients where IIV31 is found (blue) and patients where IIV31 is not found (red). Two-sided Wilcoxon rank-sum p-value is reported. **(d)** Box plots comparing CIBERSORT-inferred proportions of regulatory T cells (Tregs) and CD8 T cells between patients positive and negative for IIV31. Boxes show the quartiles (0.25 and 0.75) of the data, center lines show the medians, and whiskers show the rest of the distribution except for outliers. Source data are provided as Source Data 2. Two-sided Wilcoxon rank-sum p-value is reported for comparisons assigned with FDR q<0.05. **(e)** *Trichomonas vaginalis* and somatic mutations in *PTEN, CTNNB1* and *PIK3R1* are associated with IIV31 presence. One-sided Fisher’s exact test p-values are provided. **(f)** Bar plot comparing the fold change (relative to GAPDH) between the COV318 cell line that was predicted as *Geobacillus-positive*, and the OVISE cell line that was used as control. Error bars show the standard deviation. The one-sided t-test p-value is provided. Source data are provided as a Source Data file 4.

Notably, we identified expression in five head and neck carcinoma samples of a *Vientovirus*, a member of the recently characterized human virus family *Redondoviridae* that is associated with human oro-respiratory tract^53^ (Figure 4a, Supplementary Data 3,9). We also found expression of a *Gemycircularvirus* HV-GcV1^54^ in distinct samples from several cancer types, and *Cutavirus* expression in one COAD and one CESC sample each. We additionally detected human coxsackievirus^55^ in a COAD sample, confirming a previous report^10^.

We also found expression of a few arthropod viruses in TCGA, almost exclusively in UCEC samples (Figure 4a), most notable of which is *Armadillidium vulgare* iridescent virus (IIV31)^56^. We detected reads that align to IIV31 proteins in 152 endometrial cancer samples (which constitute more than 25% of endometrial cancer samples studied). While we did not find previous reports of IIV31 in these samples, reads that align to the same strain were recently detected in a few DNA sequencing samples, but were filtered because these were not included in databases of multiple pipelines^12^. IIV31 is in *Betairidovirinae;* members of this subfamily of dsDNA viruses infect a wide variety of arthropods, including common insect parasites of humans^57^. One study speculated on the role of *Betairidovirinae* transmitted by mosquitos in human disease^58^, but, to our knowledge, their presence in humans has not been reported before. While *Betairidovirinae* are not considered to be pathogens of vertebrates, one study showed that the model *Betairidovirinae* insect iridovirus 6 (IIV6) was lethal to mice after injection, while heat-inactivated IIV6 was not^59^. Additional studies have shown that *Betairidovirinae* can infect vertebrate predators of infected insects as well as several vertebrate cell lines^60^. Therefore, *Betairidovirinae* may opportunistically infect vertebrates, including humans.

We identified different IIV31 genes expressed in UCEC samples, and samples positive for IIV31 proteins originate from different batches and sequencing centers (Supplementary Data 10). In addition, we found that IIV31 presence was strongly and positively associated with overall survival (Figure 4b), and negatively associated with TMB and chromosome-level aneuploidy (Figure 4c, d). We did not identify a path to contamination by IIV31; the multiple origins of IIV31-positive samples and significant associations between IIV31 expression and other cancer properties both suggest that IIV31 is not a contaminant. Of the most highly expressed IIV31 proteins, we found an IAP apoptosis inhibitor homolog and serine/threonine protein kinases that were individually associated with poor overall survival (YP_009046765, YP_009046752 and YP_009046774, respectively), as well as a *RAD50* homolog (YP_009046808, Supplementary Figure 3, Supplementary Data 10).

We found significant positive association between IIV31 and CIBERSORT^61^ inferred CD8^+^ T cell frequency and Treg frequency (Figure 4d). These findings, together with the association with improved survival suggest that IIV31 could be linked with a different infection, either directly or indirectly. We explored the association of IIV31 infection with *Trichomonas vaginalis* (TV)^62^ infection. TV is a single-celled protozoan pathogen that infects the human urogenital tract^63^, and has been associated with increased risk of cervical cancer, which is enhanced by HPV coinfection^64^. We found that TV is expressed in multiple UCEC tumor samples (we verified 21 TV positive tumors with strict alignment parameters, due to high false positive rate when aligning against TV transcripts). Indeed, TV positive samples are highly enriched with IIV31 positive samples (Fisher exact test p-value = 1.4e-8). Both TV and IIV31 are significantly associated with somatic *PTEN* mutations, which are linked to better survival in endometrial cancers^65^ (whereas presence of IIV31 is also associated with mutations in *CTNNB1* and *PIK3R1*, Figure 4e).

We additionally identified *Geobacillus* virus E2 expression in 33 ovarian cancer samples; this virus is likely the most frequently expressed virus in high grade serous ovarian cancer. To further validate the presence of the *Geobacillus* virus E2, we applied viRNAtrap to cell line data from CCLE^66^. We identified the COV318 cell line as *Geobacillus* virus E2-positive and identified the OVISE cell line as a virus-negative control. Through qRT-PCR we validated the expression E2 in the predicted-positive cell line COV318 (Figure 4f). These results verify that *Geobacillus* virus E2, which was never found in ovarian cancer before, is indeed expressed in ovarian cancer cells, and that viRNAtrap can be used to sensitively detect virus-positive samples. *Geobacillus* bacteria has been previously detected in multiple ovarian cancer samples^67,68^. While we could not pinpoint the *Geobacillus* species harboring the phage, it is likely within those previously found in ovarian cancer samples^67,68^.

We found murine leukemia virus^69^ expression in distinct samples from five cancer types. However, murine leukemia virus contamination has been reported for cell culture due to human DNA preparation^70^. Our method additionally detected a previously unknown virus in a matched non-cancer sample from one HNSC patient, with protein similarity to *Pteropus* (fruit bat)-associated *Gemycircularvirus* and several other gemycircularviruses (Supplementary Data 3,9).

## Discussion

Identification of viruses from tumor RNA sequencing allows for the potential discovery of new carcinogenic agents and mechanisms. Discovery of novel and divergent viral species that contribute to cancer initiation and progression is crucial for development of new therapeutics, including vaccinations, screening practices, and antimicrobial treatments. Viruses are currently identified from sequencing reads based on similarity to known viruses^71^. However, when studying viruses from short reads, typical with Illumina-based RNA sequencing, reads originating from divergent viruses may share little sequence similarity to known viruses, rendering the identification of novel viruses highly challenging.

To address this challenge, we developed viRNAtrap, a new, alignment-free framework to identify viral reads from RNAseq and assemble viral contigs. The contigs detected by viRNAtrap can be aligned to different viral databases, as we demonstrate in this study, to rapidly identify viral expression of interest in tumor samples. We curate a database of HERVs that comprise intact retroviral genes in the human genome and survey the expression of these viruses across different cancer tissues. Through a database of divergent viruses, we demonstrate that viRNAtrap identifies viruses in TCGA samples that were not detected in previous studies. This is enabled through an integrative method that uses the model scores to assemble viral reads rather than aligning short divergent reads to viral databases or applying de-novo assembly to many unmapped reads. We further show that using the deep learning model substantially improves the running time, while not compromising sensitivity if more than 5 viral reads are present (Supplementary Figure 4, see Methods). Importantly, the output of viRNAtrap can be alternatively used as input to motif search tools, to potentially identify highly divergent viruses. Because the deep learning model underlying viRNAtrap was trained to distinguish viral from human sequences, the model predictions for sequences derived from a range of other organisms is not defined. Future work could train models to identify viruses from a variety of other organisms, and, with the viRNAtrap framework, achieve higher sensitivity for viral detection.

We employ viRNAtrap for an exploratory data analysis and characterize viruses that are expressed across 14 cancer tissues from TCGA and analyze their genomic and survival correlates. Interestingly, while the expression of some exogenous cancer viruses is known to be associated with improved survival, we found that the expression of human endogenous viruses is strictly associated with poor survival rates. Expression of a virus of the subfamily *Betairidovirinae*, which are pathogens of insects, found in endometrial cancer tissues was similarly associated with significantly better overall patient survival. For all divergent viruses reported in this study, the presence and classification of multiple viral reads was verified by targeted blastn- and blastx-based sequence analyses in different samples. However, it is not possible to model all contaminant of viruses that may have infected the samples during laboratory procedures^16^.

Perhaps, the most interesting divergent virus we found is IIV31 from the subfamily *Betairidoovirinae*, which was frequently detected in UCEC TCGA samples. Interestingly, IIV6, a very close relative of IIV31, can infect a variety of vertebrates including mice, and induces an immune response in mammalian tissues^60,72^. Thus, one possibility is that IIV31 is transmitted to the uterus through another insect, such as the crab louse. While we have not yet confirmed the source of this virus, our results imply that its presence may be a direct or indirect consequence of *Trichomonas vaginalis* infection. Therefore, it shows that viRNAtrap is sufficiently powerful to identify a previously unknown viral transcript in tumor samples, whether oncogenic or neutral. Through this analysis, we also identified TV reads in multiple endometrial cancer samples, indicating a possible new association between TV and endometrial cancer, like the known association of TV with cervical cancer^64^. One of the established pathogenic mechanisms of TV infection in humans, which may also explain the frequent HPV coinfection, is that TV secretes exosomes that have the effect of suppressing CXCL8^73^. Interestingly, low expression of CXCL8, like infection with TV, has been associated with favorable prognosis in cervical cancer^74^. Thus, it is possible that the presence of IIV31 is a secondary infection in patients already infected with TV or some other pathogen that suppresses the human anti-viral response.

Importantly, we identified *E2 Geobacillus* virus in 10% of high-grade, serous ovarian cancers, making it the most frequently expressed virus in this cancer type. We experimentally verified that *E2 Geobacillus* is indeed expressed in cell lines. We also found expression of a *Redondoviridae* member in head and neck cancers that was not previously reported^75^. This finding calls for a study of the role of *Redondoviridae* in tumor initiation and progression, as this family of viruses was only recently detected in humans and associated with different clinical conditions.

In conclusion, we developed viRNAtrap, a new software for alignment free identification of viruses from RNAseq, allowing rapid characterization of viral expression and detection of divergent viruses. We applied it to tumor tissues from TCGA, uncovering expression patterns of different groups of viruses. We report previously unrecognized associations between several forms of cancer and several unexpected viral clades, including viral clades canonically found in produce and in insect parasites of humans. Future studies may employ viRNAtrap to find viruses that contribute to other malignancies.

## Methods

### Training a neural network to distinguish viral RNA sequencing reads

The viRNAtrap framework is composed of two main components, illustrated in figure 1a. The first is a deep learning model, which was trained to accurately distinguish viral from human reads using RNA-sequencing. The second assembles the predicted viral reads into contigs. The trained neural network is composed of one 1D-convolutional layer and three fully connected layers, one of which is the final output layer. The RNA sequences were one-hot encoded to vectors that were given as input to the model. The learning rate was set to 0.0005, we used 64 filters with ReLU as an activation function in the convolutional layer, followed by one pooling layer for feature extraction. The global extracted features from the convolutional layer are passed to three fully connected layers, to make a prediction based on a sigmoid activation function in the output layer.

To train the model, we collected human and viral sequencing data. Coding sequences of human and other placentals viruses downloaded from the Virus Variation Resource^76^. Human transcripts for hg19 were downloaded from NCBI Human Genome Resources^77^. These sequences were segmented into 48bp segments, which is the read length for the RNAseq in almost all tumor types in TCGA; only a few tumor types that were added chronologically last to TCGA used longer reads. We used a 48bp window size for human transcripts and a 2bp window size for viral sequences, to balance the positive and negative data. Then, these were randomly split (where all segments of each transcript were considered together) into balanced train, validation, and test sets (n= 8,000,000, 800,000, and 2,558,044, respectively).

### Model performance evaluation and comparison to existing methods

We evaluated the performance of the model using the Area Under the Receiver Operating Characteristic Curve (AUROC), the Area Under the Precision Recall Curve (AUPRC), as well as accuracy, precision, recall, and F1-score, for the test dataset. We trained multiple models with different architectures and hyperparameters and then selected the model with highest average between the validation-set AUROC and recall. The model was trained using TensorFlow 2.6.0 and Keras^78^. We compared the performance of our model to models from DeepViFi^16^, DeepVirFinder^15^, ViraMiner^21^ and off-the-shelf Seq2Seq model. Because this is the first approach trained to predict viruses from RNA sequencing reads of length 48bp, we used our training data to retrain each of these models, following the instructions provided by each method, and evaluated the AUROC, AUPRC, accuracy, precision, recall, and F1-score using our test set (see Supplementary Methods for detailed description of hyper-parameters used). Importantly, existing methods were not designed for reads shorted than 150bp, therefore they should not be expected to perform as well as viRNAtrap on 48bp segments, for which viRNAtrap was optimized. Our comparison does not rule out the possibility that new hyper parameter optimization for this purpose may enhance the performance of existing methods for 48bp sequences.

### Assembling viral contigs from neural network predicted viral reads

Once the viRNAtrap model predicts the probability of a viral origin of each read, reads with model scores more than 0.7 are used as seeds to assemble viral contigs. Viral contigs are assembled using iterative search for substrings with exact matches between 24bp k-mers. Each seed is complemented from the left and right ends using its left-most and right-most 24bp k-mers. For both the left and right assembly, reads containing the left or right most k-mers in a different position from the read that is being searched are identified. The read adding the maximal number of bases to the assembled contig is used to complement the left and right contigs. The model scores that were assigned to reads that are used to assemble each contig were averaged, and the assembly terminates if the average score is below 0.5. Finally, the right and left contigs are concatenated, to yield a complete viral contig. This algorithm was implemented in Python 3 and subsequently in C, which improved the running time by more than an order of magnitude for inputs with large numbers of reads.

### Data pre-processing

We downloaded RNA-sequencing data from Genomic Data Commons (GDC; https://portal.gdc.cancer.gov/)79 as BAM files. High quality reads were selected and mapped with Bowtie2 against hg19 (1000 Genomes version) and PhiX phage (NC_001422), and only the unmapped reads were kept. Then, we merged the paired end reads and converted them to fastq files, which were used as input to for the viRNAtrap framework, to yield predicted viral contigs.

### Viral databases

Viral contigs yielded by the assembly component were used as inputs to blastn^24^. Three databases were used to search for viruses (with E-value threshold of 0.01):

(1) RefSeq reference human viruses, downloaded from the National Center for Biotechnology Information (NCBI) ^77^, to which we added human papillomaviruses strains that are not in RefSeq from PAVE (https://pave.niaid.nih.gov)^80^. Reference viruses were searched using blastn, with default parameters except for a word size of 15 (lower than the default of 28), which was chosen to allow identification from short contigs.

(2) more divergent viruses obtained from RVDB^81^ (https://hive.biochemistry.gwu.edu/rvdb/) which was then filtered to remove non-viral elements, endogenous viruses, and accessions that were consistently not verified using blastn against the nonredundant (nr) blast nucleotide database.

(3) Human endogenous viruses. We curated a database of potentially functional HERVs through evaluation of viral protein completeness (in contrast to a previous study that evaluated HERV expression in distinct RNAseq datasets^82^). The initial genomic locations of reported HERV elements were downloaded from the HERVd HERV annotation database (https://herv.img.cas.cz)^83^. The nucleotide sequences in hg19 for each reported HERV were extracted using twoBitToFa^84^. We then applied blastx against NR with E-value cutoff of 1E-4, as well as a profile search^85^ against collected POL proteins, where the profile was obtained by collecting POL genes annotated in GenBank in lentiviruses (as of September 2016) and aligning their amino acid sequences using MAFFT^86^. Sequences with at least one identified retroviral protein motif of: POL/RT, GAG or ENV were extracted, yielding 3,044 HERVs that were considered for search in TCGA samples (Supplementary Data 5). Importantly, high mutation rate of HERV^87^ prohibits most HERV sequences from aligning to the human genome in pre-processing^12,88^, however, in rare cases, HERV regions that are conserved would not be identified by this approach.

### Quality standards for virus identification

For all viruses, blastn was applied with E-value cutoff of 0.01 and any sequences with a match to contaminant accessions (that were associated with vector contamination) were filtered out.

a. Reference viruses. For every sample, contigs mapped to each accession were extracted. Identified accessions with maximum qcov across contigs more that 90%, average qcov more than 50%, and average similarity more than 90% were considered. Accessions with maximal contig length under 100bp were manually inspected and verified against nr.
b. Human endogenous viruses. For every sample, contigs mapped to each HERV were extracted. HERVs with contigs longer that 200bp, and with average qcov and similarity more than 95% were considered.
c. Divergent viruses. For every sample, contigs mapped to each accession were extracted. Viruses already identified through the reference database were removed. Identified accessions with maximal contig length more than 300bp and qcov more than 40%, or with maximal contig length more than 100bp and qcov more than 75% and average similarity more than 75% were considered for manual inspection.

All instances of divergent viruses identified in TCGA samples were verified using blastn against nr, to support that the virus strain is indeed the best match to a viral contig generated by viRNAtrap. We reason that non-reference viruses (divergent viruses and viruses of nonhuman hosts) that were identified and verified in more than one sample were less likely to be contaminant or isolated events, whereas sample with fewer reads from such viruses may be filtered due to the strict filtering. We therefore additionally searched using the STAR aligner^89^ across tumor types where these viruses were identified through viRNAtrap (Supplementary Data 3). The following accessions were additionally searched using STAR to increase sample coverage (as these were the most interesting divergent strains found across multiple samples): Bermuda grass latent virus (NC_032405), *Armadillidium vulgare* iridescent virus IIV31 (NC_024451), *Geobacillus* virus (NC_009552) and the Human lung-associated vientovirus (NC_055523)

### Filtering contaminants

To filter vector contaminants, we applied VeScreen^90^ to the assembled contigs that have been mapped to viruses through our databases, where virus accessions associated with vector contaminants were entirely removed from the search (Supplementary Data 11).

In addition, we examined the application of software such as Kraken2^91^ to the RNAseq reads for filtering reads that are not likely of viral origin, by applying Kraken2 to reads of LIHC samples. However, we found that 99% of the reads would not be filtered using this approach (Supplementary Figure 5), likely due to the short reads (48bp) for which Kraken has not been designed or evaluated, as longer sequences are known to be more accurately mapped^92^.

### Genomic correlates of viral expression

We correlated viral expression with genomic markers across TCGA samples. Chromosomal aneuploidy levels for TCGA samples were extracted from^93^ and the total number of chromosome-arm-level alterations was used. The tumor mutation burden was defined to be the total number of somatic mutations in each sample, downloaded from the Xena browser^94^ (https://xenabrowser.net). CIBERSORT^61^ software was applied to TCGA samples using the default set of 22 immune-cell signatures.

### Cells and culture conditions

Human ovarian cancer cell lines COV318 and OVISE were cultured in RPMI1640 medium containing 10% fetal bovine serum (FBS) and 1% penicillin-streptomycin under 5% CO_2_. All of the cell lines were authenticated at The Wistar Institute’s Genomics Facility using shorttandem-repeat DNA profiling. Regular mycoplasma testing was performed using a LookOut mycoplasma PCR detection kit (Sigma, cat. no. MP0035).

### Experimental validation of the *Geobacillus* virus E2 in ovarian cancer cell lines

Reverse-transcriptase qPCR (RT-qPCR) RNA was extracted using TRIzol reagent (Invitrogen, cat. no. 15596026). Extracted RNA was used for reverse-transcriptase PCR using a High-capacity cDNA reverse transcription kit (Thermo Fisher, cat. no. 4368814). Quantitative PCR was performed using a QuantStudio 3 real-time PCR system. GAPDH was used as an internal control. The fold change was calculated using the 2-ΔΔCt method. The primers used for reverse-transcriptase qPCR are: GAPDH forward, GTCTCCTCTGACTTCAACAGCG and reverse, ACCACCCTGTTGCTGTAGTAGCCAA; *Geobacillus* virus E2 terminase forward, TTGCGATGCGTACTCAGACT and reverse, CTCTTTTTGGTCAGCAGCGG Primers were obtained using NCBI primer design tool as shown in the attached word document. The primers were synthesized by Integrated DNA Technologies IDT. The document specifying the primer design is provided in the Supplementary Information.

### Identification of *Trichomonas vaginalis-positive* samples

UCEC unmapped (to hg19) reads were aligned to the reference genome of *Trichomonas vaginalis (GCF_000002825)^62^* strain G3 using blastn^24^ with E-value < 1e-8 and more than 90% identity. These thresholds were set to remove false positives that were frequent when aligning against *Trichomonas vaginalis* when examining both blastn^24^ and STAR aligner^89^. TV reads for each TV-positive sample were verified by manual inspection of the output alignments.

### viRNAtrap performance evaluation

To evaluate the contribution of the model to the viRNAtrap pipeline we re-ran viRNAtrap on 10 LIHC samples, and additionally ran a modified viRNAtrap pipeline not using the model, on the same system. We compared the viruses identified by the model-based approach to those that are identified when the pipeline is applied without using the model (Supplementary Table 2, showing similar viruses with different number of contigs). We additionally compared the running time of the two approaches (Supplementary Figure 4).

To evaluate the sensitivity of the viRNAtrap pipeline based the number of viral reads present in a sample, we performed a simulated analysis. From the test dataset, we down sampled groups of viral reads with different group sizes (10,000 groups for each size, from one read up to 10 reads), and we evaluated the number of groups with at least one read that is scored above 0.7, which is the seed threshold used for the viRNAtrap assembly. Therefore, this analysis is estimating the probability of identifying viruses based on the number of reads present. We found 93% and 99% of the groups with more than 5 and 9 reads, respectively, would be identified.

### Statistical methods

Survival analysis, including Kaplan Meier curves plots, log rank test p-values were obtained using the Python lifelines package (v0.26.4)^95^. P-values comparing TMB and aneuploidy between two groups correspond were computed with two-sided Wilcoxon rank-sum tests. Heatmap clustograms were generated through seaborn clustermap.

Viruses with significant log-rank p-values are reported as significantly associated with survival.

None of the reference viruses were significantly associated with survival after FDR correction (Supplementary Table 1), however, we report in Figure 2 the association between HR-HPV with unadjusted p-value because it is confirmatory of a known association between HR-HPV and HNSC survival^33,34^.

For HERV, our exploratory data analysis uncovered some significant associations with the complete hypothesis testing. We present in the main text selected associations with at least 5 cases in each group. Nevertheless, FDR correction was applied within each cancer type for all HERV associations, and we additionally applied a global FDR correction for all comparisons across cancer types, yielding some significant associations with less than 5 positive cases. The complete significant associations between survival and viral presence are reported in the Supplementary Data 12.

## Data Availability

The complete training and test data as well as viral databases generated in this study have been deposited in the Zenodo database under accession code https://doi.org/10.5281/zenodo.7548375. The results shown here are in whole or part based upon data generated by the TCGA Research Network: https://www.cancer.gov/tcga. The raw FASTQ RNA sequencing data are protected and are not publicly available due to data privacy laws, but are available under restricted access as data can be unique to an individual. Access can be obtained from the Genome Data Commons (GDC) after receiving permission via dbGaP, following the steps described in: https://www.ncbi.nlm.nih.gov/projects/gap/cgi-bin/study.cgi?studyid=phs000178.v11.p8. The processed data including viruses identified and respective statistics are available as supplementary Data 3. The complete data generated in this study are provided in the Supplementary Information/Source Data file. Source data are provided with this paper.

## Code Availability

The scripts for pre and post processing and the viRNAtrap package are available through GitHub: https://github.com/AuslanderLab/virnatrap and Zenodo under accession code: https://doi.org/10.5281/zenodo.7548375.

## Acknowledgements

The research reported in this publication was supported in part by the National Cancer Institute of the National Institutes of Health under Award Number R00CA252025 (N.A.), RF1-AG063481, P30-CA016520 (Q.L.), NIH RO1 AI15350 (P.M.L) Commonwealth of Pennsylvania SAP# 4100089371, and P30 CA010815, and by the Intramural Research Program of the National Institutes of Health, National Cancer Institute (A.A.S).

## Author Contributions Statement

N.A. initiated the project. Q.L., R.Z., A.A.S. and N.A. supervised work. A.E., X.H., R.Z., A.A.S. and N.A. designed and performed experiments and analyses. A.E., Y.Y., D.E.S., J.W., A.A.S. and N.A wrote and tested software. P.M.L and Q.M. contributed to data interpretation and exploratory analyses. K.T. and Q.L. revised the survival analysis.

## References

1 Morales-Sánchez, A. & Fuentes-Pananá, E. M. Human viruses and cancer. Viruses 6, 4047–4079, doi:10.3390/v6104047 (2014).

2 Krump, N. A. & You, J. Molecular mechanisms of viral oncogenesis in humans. Nat Rev Microbiol 16, 684–698, doi:10.1038/s41579-018-0064-6 (2018).

3 Rous, P. A SARCOMA OF THE FOWL TRANSMISSIBLE BY AN AGENT SEPARABLE FROM THE TUMOR CELLS. J Exp Med 13, 397–411, doi:10.1084/jem.13.4.397 (1911).

4 Moore, P. S. & Chang, Y. Why do viruses cause cancer? Highlights of the first century of human tumour virology. Nat Rev Cancer 10, 878–889, doi:10.1038/nrc2961 (2010).

5 Salyakina, D. & Tsinoremas, N. F. Viral expression associated with gastrointestinal adenocarcinomas in TCGA high-throughput sequencing data. Hum Genomics 7, 23, doi:10.1186/1479-7364-7-23 (2013).

6 Parfenov, M. et al. Characterization of HPV and host genome interactions in primary head and neck cancers. Proc Natl Acad Sci U S A 111, 15544–15549, doi:10.1073/pnas.1416074111 (2014).

7 Cao, S. et al. Divergent viral presentation among human tumors and adjacent normal tissues. Sci Rep 6, 28294, doi:10.1038/srep28294 (2016).

8 Strong, M. J. et al. Differences in gastric carcinoma microenvironment stratify according to EBV infection intensity: implications for possible immune adjuvant therapy. PLoS Pathog 9, e1003341, doi:10.1371/journal.ppat.1003341 (2013).

9 Khoury, J. D. et al. Landscape of DNA virus associations across human malignant cancers: analysis of 3,775 cases using RNA-Seq. J Virol 87, 8916–8926, doi:10.1128/jvi.00340-13 (2013).

10 Tang, K. W., Alaei-Mahabadi, B., Samuelsson, T., Lindh, M. & Larsson, E. The landscape of viral expression and host gene fusion and adaptation in human cancer. Nat Commun 4, 2513, doi:10.1038/ncomms3513 (2013).

11 Cantalupo, P. G., Katz, J. P. & Pipas, J. M. Viral sequences in human cancer. Virology 513, 208–216, doi:10.1016/j.virol.2017.10.017 (2018).

12 Zapatka, M. et al. The landscape of viral associations in human cancers. Nat Genet 52, 320–330, doi:10.1038/s41588-019-0558-9 (2020).

13 Kellam, P. Molecular identification of novel viruses. Trends Microbiol 6, 160–165, doi:10.1016/s0966-842x(98)01239-6 (1998).

14 Ren, J., Ahlgren, N. A., Lu, Y. Y., Fuhrman, J. A. & Sun, F. VirFinder: a novel k-mer based tool for identifying viral sequences from assembled metagenomic data. Microbiome 5, 69, doi:10.1186/s40168-017-0283-5 (2017).

15 Ren, J. et al. Identifying viruses from metagenomic data using deep learning. Quant Biol 8, 64–77, doi:10.1007/s40484-019-0187-4 (2020).

16 Rajkumar, U. et al. in Proceedings of the 13th ACM International Conference on Bioinformatics, Computational Biology and Health Informatics 1–8 (2022).

17 Fang, Z. et al. PPR-Meta: a tool for identifying phages and plasmids from metagenomic fragments using deep learning. Gigascience 8, doi:10.1093/gigascience/giz066 (2019).

18 Auslander, N., Gussow, A. B., Benler, S., Wolf, Y. I. & Koonin, E. V. Seeker: alignment-free identification of bacteriophage genomes by deep learning. Nucleic Acids Res 48, e121, doi:10.1093/nar/gkaa856 (2020).

19 Zhang, Z. et al. Rapid identification of human-infecting viruses. Transbound Emerg Dis 66, 2517–2522, doi:10.1111/tbed.13314 (2019).

20 Bartoszewicz, J. M., Seidel, A. & Renard, B. Y. Interpretable detection of novel human viruses from genome sequencing data. NAR Genom Bioinform 3, lqab004, doi:10.1093/nargab/lqab004 (2021).

21 Tampuu, A., Bzhalava, Z., Dillner, J. & Vicente, R. ViraMiner: Deep learning on raw DNA sequences for identifying viral genomes in human samples. PLoS One 14, e0222271, doi:10.1371/journal.pone.0222271 (2019).

22 Weinstein, J. N. et al. The Cancer Genome Atlas Pan-Cancer analysis project. Nat Genet 45, 1113–1120, doi:10.1038/ng.2764 (2013).

23 Mukherjee, S., Huntemann, M., Ivanova, N., Kyrpides, N. C. & Pati, A. Large-scale contamination of microbial isolate genomes by Illumina PhiX control. Stand Genomic Sci 10, 18, doi:10.1186/1944-3277-10-18 (2015).

24 Altschul, S. F. et al. Gapped BLAST and PSI-BLAST: a new generation of protein database search programs. Nucleic Acids Res 25, 3389–3402, doi:10.1093/nar/25.17.3389 (1997).

25 Coursey, T. L., Van Doorslaer, K. & McBride, A. A. Regulation of Human Papillomavirus 18 Genome Replication, Establishment, and Persistence by Sequences in the Viral Upstream Regulatory Region. J Virol 95, e0068621, doi:10.1128/jvi.00686-21 (2021).

26 Doorbar, J. et al. The biology and life-cycle of human papillomaviruses. Vaccine 30 Suppl 5, F55–70, doi:10.1016/j.vaccine.2012.06.083 (2012).

27 Schaffer, A. A. et al. Integration of adeno-associated virus (AAV) into the genomes of most Thai and Mongolian liver cancer patients does not induce oncogenesis. BMC Genomics 22, 814, doi:10.1186/s12864-021-08098-9 (2021).

28 Bayard, Q. et al. Cyclin A2/E1 activation defines a hepatocellular carcinoma subclass with a rearrangement signature of replication stress. Nat Commun 9, 5235, doi:10.1038/s41467-018-07552-9 (2018).

29 Cossart, Y. E., Field, A. M., Cant, B. & Widdows, D. Parvovirus-like particles in human sera. Lancet 1, 72–73, doi:10.1016/s0140-6736(75)91074-0 (1975).

30 Adamson-Small, L. A., Ignatovich, I. V., Laemmerhirt, M. G. & Hobbs, J. A. Persistent parvovirus B19 infection in non-erythroid tissues: possible role in the inflammatory and disease process. Virus Res 190, 8–16, doi:10.1016/j.virusres.2014.06.017 (2014).

31 Dickinson, A. et al. Newly detected DNA viruses in juvenile nasopharyngeal angiofibroma (JNA) and oral and oropharyngeal squamous cell carcinoma (OSCC/OPSCC). Eur Arch Otorhinolaryngol 276, 613–617, doi:10.1007/s00405-018-5250-7 (2019).

32 Li, Y. et al. Detection of parvovirus B19 nucleic acids and expression of viral VP1/VP2 antigen in human colon carcinoma. Am J Gastroenterol 102, 1489–1498, doi:10.1111/j.1572-0241.2007.01240.x (2007).

33 Sethi, S. et al. Characteristics and survival of head and neck cancer by HPV status: a cancer registry-based study. Int J Cancer 131, 1179–1186, doi:10.1002/ijc.26500 (2012).

34 Sarkar, S. et al. Human papilloma virus (HPV) infection leads to the development of head and neck lesions but offers better prognosis in malignant Indian patients. Med Microbiol Immunol 206, 267–276, doi:10.1007/s00430-017-0502-5 (2017).

35 Curty, G. et al. Human Endogenous Retrovirus K in Cancer: A Potential Biomarker and Immunotherapeutic Target. Viruses 12, doi:10.3390/v12070726 (2020).

36 Kolbe, A. R. et al. Human Endogenous Retrovirus Expression Is Associated with Head and Neck Cancer and Differential Survival. Viruses 12, doi:10.3390/v12090956 (2020).

37 Kammerer, U., Germeyer, A., Stengel, S., Kapp, M. & Denner, J. Human endogenous retrovirus K (HERV-K) is expressed in villous and extravillous cytotrophoblast cells of the human placenta. J Reprod Immunol 91, 1–8, doi:10.1016/j.jri.2011.06.102 (2011).

38 Armbruester, V. et al. A novel gene from the human endogenous retrovirus K expressed in transformed cells. Clin Cancer Res 8, 1800–1807 (2002).

39 Wang-Johanning, F. et al. Human endogenous retrovirus K triggers an antigen-specific immune response in breast cancer patients. Cancer Res 68, 5869–5877, doi:10.1158/0008-5472.Can-07-6838 (2008).

40 Wang-Johanning, F. et al. Expression of human endogenous retrovirus k envelope transcripts in human breast cancer. Clin Cancer Res 7, 1553–1560 (2001).

41 Kassiotis, G. Endogenous retroviruses and the development of cancer. J Immunol 192, 1343–1349, doi:10.4049/jimmunol.1302972 (2014).

42 Xue, B., Sechi, L. A. & Kelvin, D. J. Human Endogenous Retrovirus K (HML-2) in Health and Disease. Front Microbiol 11, 1690, doi:10.3389/fmicb.2020.01690 (2020).

43 Kim, J. S., Yoon, S. J., Park, Y. J., Kim, S. Y. & Ryu, C. M. Crossing the kingdom border: Human diseases caused by plant pathogens. Environ Microbiol 22, 2485–2495, doi:10.1111/1462-2920.15028 (2020).

44 Hahn, S. et al. Serological response to human endogenous retrovirus K in melanoma patients correlates with survival probability. AIDS Res Hum Retroviruses 24, 717–723, doi:10.1089/aid.2007.0286 (2008).

45 Zhao, J. et al. Expression of Human Endogenous Retrovirus Type K Envelope Protein is a Novel Candidate Prognostic Marker for Human Breast Cancer. Genes Cancer 2, 914–922, doi:10.1177/1947601911431841 (2011).

46 Reis, B. S. et al. Prostate cancer progression correlates with increased humoral immune response to a human endogenous retrovirus GAG protein. Clin Cancer Res 19, 6112–6125, doi:10.1158/1078-0432.Ccr-12-3580 (2013).

47 Fan, X., Quezada, S. A., Sepulveda, M. A., Sharma, P. & Allison, J. P. Engagement of the ICOS pathway markedly enhances efficacy of CTLA-4 blockade in cancer immunotherapy. J Exp Med 211, 715–725, doi:10.1084/jem.20130590 (2014).

48 Xiao, Z., Mayer, A. T., Nobashi, T. W. & Gambhir, S. S. ICOS Is an Indicator of T-cell-Mediated Response to Cancer Immunotherapy. Cancer Res 80, 3023–3032, doi:10.1158/0008-5472.Can-19-3265 (2020).

49 Faget, J. et al. ICOS-ligand expression on plasmacytoid dendritic cells supports breast cancer progression by promoting the accumulation of immunosuppressive CD4+ T cells. Cancer Res 72, 6130–6141, doi:10.1158/0008-5472.Can-12-2409 (2012).

50 Conrad, C. et al. Plasmacytoid dendritic cells promote immunosuppression in ovarian cancer via ICOS costimulation of Foxp3(+) T-regulatory cells. Cancer Res 72, 5240–5249, doi:10.1158/0008-5472.Can-12-2271 (2012).

51 Zhang, T. et al. RNA viral community in human feces: prevalence of plant pathogenic viruses. PLoS Biol 4, e3, doi:10.1371/journal.pbio.0040003 (2006).

52 Balique, F., Lecoq, H., Raoult, D. & Colson, P. Can plant viruses cross the kingdom border and be pathogenic to humans? Viruses 7, 2074–2098, doi:10.3390/v7042074 (2015).

53 Abbas, A. A. et al. Redondoviridae, a Family of Small, Circular DNA Viruses of the Human Oro-Respiratory Tract Associated with Periodontitis and Critical Illness. Cell Host Microbe 25, 719–729.e714, doi:10.1016/j.chom.2019.04.001 (2019).

54 Halary, S. et al. Novel Single-Stranded DNA Circular Viruses in Pericardial Fluid of Patient with Recurrent Pericarditis. Emerg Infect Dis 22, 1839-1841, doi:10.3201/eid2210.160052 (2016).

55 Dalldorf, G. & Sickles, G. M. An Unidentified, Filtrable Agent Isolated From the Feces of Children With Paralysis. Science 108, 61–62, doi:10.1126/science.108.2794.61 (1948).

56 Federici, B. A. Isolation of an iridovirus from two terrestrial isopods, the pill bug, Armadillidium vulgare, and the sow bug, Porcellio dilatatus. Journal of Invertebrate Pathology 36, 373–381, doi:https://doi.org/10.1016/0022-2011(80)90041-5 (1980).

57 Williams, T. Natural invertebrate hosts of iridoviruses (Iridoviridae). Neotrop Entomol 37, 615–632, doi:10.1590/s1519-566x2008000600001 (2008).

58 Li, L. et al. Investigation on Mosquito-Borne Viruses at Lancang River and Nu River Watersheds in Southwestern China. Vector Borne Zoonotic Dis 17, 804–812, doi:10.1089/vbz.2017.2164 (2017).

59 Ohba, M. & Aizawa, K. Mammalian toxicity of an insect iridovirus. Acta Virol 26, 165–168 (1982).

60 ince i, A., Özcan, O., Ilter-Akulke, A. Z., Scully, E. D. & Özgen, A. Invertebrate Iridoviruses: A Glance over the Last Decade. Viruses 10, doi:10.3390/v10040161 (2018).

61 Newman, A. M. et al. Robust enumeration of cell subsets from tissue expression profiles. Nat Methods 12, 453–457, doi:10.1038/nmeth.3337 (2015).

62 Carlton, J. M. et al. Draft genome sequence of the sexually transmitted pathogen Trichomonas vaginalis. Science 315, 207–212, doi:10.1126/science.1132894 (2007).

63 Kissinger, P. Trichomonas vaginalis: a review of epidemiologic, clinical and treatment issues. BMC Infect Dis 15, 307, doi:10.1186/s12879-015-1055-0 (2015).

64 Yang, S. et al. Trichomonas vaginalis infection-associated risk of cervical cancer: A meta-analysis. Eur J Obstet Gynecol Reprod Biol 228, 166–173, doi:10.1016/j.ejogrb.2018.06.031 (2018).

65 Risinger, J. I. et al. PTEN mutation in endometrial cancers is associated with favorable clinical and pathologic characteristics. Clin Cancer Res 4, 3005–3010 (1998).

66 Barretina, J. et al. The Cancer Cell Line Encyclopedia enables predictive modelling of anticancer drug sensitivity. Nature 483, 603–607, doi:10.1038/nature11003 (2012).

67 Banerjee, S. et al. The ovarian cancer oncobiome. Oncotarget 8, 36225–36245, doi:10.18632/oncotarget.16717 (2017).

68 Nejman, D. et al. The human tumor microbiome is composed of tumor type-specific intracellular bacteria. Science 368, 973–980, doi:10.1126/science.aay9189 (2020).

69 Robinson, H. L. Retroviruses and cancer. Rev Infect Dis 4, 1015–1025, doi:10.1093/clinids/4.5.1015 (1982).

70 Uphoff, C. C., Lange, S., Denkmann, S. A., Garritsen, H. S. & Drexler, H. G. Prevalence and characterization of murine leukemia virus contamination in human cell lines. PLoS One 10, e0125622, doi:10.1371/journal.pone.0125622 (2015).

71 Kostic, A. D. et al. PathSeq: software to identify or discover microbes by deep sequencing of human tissue. Nat Biotechnol 29, 393–396, doi:10.1038/nbt.1868 (2011).

72 Ahlers, L. R., Bastos, R. G., Hiroyasu, A. & Goodman, A. G. Invertebrate Iridescent Virus 6, a DNA Virus, Stimulates a Mammalian Innate Immune Response through RIG-I-Like Receptors. PLoS One 11, e0166088, doi:10.1371/journal.pone.0166088 (2016).

73 Twu, O. et al. Trichomonas vaginalis exosomes deliver cargo to host cells and mediate host:parasite interactions. PLoS Pathog 9, e1003482, doi:10.1371/journal.ppat.1003482 (2013).

74 Wu, X. et al. Identification of Key Genes and Pathways in Cervical Cancer by Bioinformatics Analysis. Int J Med Sci 16, 800–812, doi:10.7150/ijms.34172 (2019).

75 Taylor, L. J. et al. Redondovirus Diversity and Evolution on Global, Individual, and Molecular Scales. J Virol 95, e0081721, doi:10.1128/jvi.00817-21 (2021).

76 Hatcher, E. L. et al. Virus Variation Resource - improved response to emergent viral outbreaks. Nucleic Acids Res 45, D482–d490, doi:10.1093/nar/gkw1065 (2017).

77 Sayers, E. W. et al. Database resources of the National Center for Biotechnology Information. Nucleic Acids Res 49, D10–d17, doi:10.1093/nar/gkaa892 (2021).

78 Keras (2015).

79 Grossman, R. L. et al. Toward a Shared Vision for Cancer Genomic Data. N Engl J Med 375, 1109–1112, doi:10.1056/NEJMp1607591 (2016).

80 Van Doorslaer, K. et al. The Papillomavirus Episteme: a major update to the papillomavirus sequence database. Nucleic Acids Res 45, D499–d506, doi:10.1093/nar/gkw879 (2017).

81 Goodacre, N., Aljanahi, A., Nandakumar, S., Mikailov, M. & Khan, A. S. A Reference Viral Database (RVDB) To Enhance Bioinformatics Analysis of High-Throughput Sequencing for Novel Virus Detection. mSphere 3, doi:10.1128/mSphereDirect.00069-18 (2018).

82 Tokuyama, M. et al. ERVmap analysis reveals genome-wide transcription of human endogenous retroviruses. Proc Natl Acad Sci U S A 115, 12565–12572, doi:10.1073/pnas.1814589115 (2018).

83 Paces, J. et al. HERVd: the Human Endogenous Retroviruses Database: update. Nucleic Acids Res 32, D50, doi:10.1093/nar/gkh075 (2004).

84 Karolchik, D. et al. The UCSC Table Browser data retrieval tool. Nucleic Acids Res 32, D493–496, doi:10.1093/nar/gkh103 (2004).

85 Yutin, N., Puigbò, P., Koonin, E. V. & Wolf, Y. I. Phylogenomics of prokaryotic ribosomal proteins. PLoS One 7, e36972, doi:10.1371/journal.pone.0036972 (2012).

86 Katoh, K. & Standley, D. M. MAFFT multiple sequence alignment software version 7: improvements in performance and usability. Mol Biol Evol 30, 772–780, doi:10.1093/molbev/mst010 (2013).

87 Bannert, N. & Kurth, R. Retroelements and the human genome: new perspectives on an old relation. Proc Natl Acad Sci U S A 101 Suppl 2, 14572–14579, doi:10.1073/pnas.0404838101 (2004).

88 Smith, C. C. et al. Endogenous retroviral signatures predict immunotherapy response in clear cell renal cell carcinoma. J Clin Invest 128, 4804–4820, doi:10.1172/jci121476 (2018).

89 Dobin, A. et al. STAR: ultrafast universal RNA-seq aligner. Bioinformatics 29, 15–21, doi:10.1093/bioinformatics/bts635 (2013).

90 Schaffer, A. A. et al. VecScreen_plus_taxonomy: imposing a tax(onomy) increase on vector contamination screening. Bioinformatics 34, 755–759, doi:10.1093/bioinformatics/btx669 (2018).

91 Wood, D. E., Lu, J. & Langmead, B. Improved metagenomic analysis with Kraken 2. Genome Biol 20, 257, doi:10.1186/s13059-019-1891-0 (2019).

92 Celaj, A., Markle, J., Danska, J. & Parkinson, J. Comparison of assembly algorithms for improving rate of metatranscriptomic functional annotation. Microbiome 2, 39, doi:10.1186/2049-2618-2-39 (2014).

93 Taylor, A. M. et al. Genomic and Functional Approaches to Understanding Cancer Aneuploidy. Cancer Cell 33, 676–689.e673, doi:10.1016/j.ccell.2018.03.007 (2018).

94 Goldman, M. J. et al. Visualizing and interpreting cancer genomics data via the Xena platform. Nat Biotechnol 38, 675–678, doi:10.1038/s41587-020-0546-8 (2020).

95 Davidson-Pilon, C. lifelines: survival analysis in Python. Journal of Open Source Software 4, 1317, doi:10.21105/joss.01317 (2019).

